# The Human walking kinematic model combined with upper and lower limb movement

**DOI:** 10.1101/2021.02.06.430092

**Authors:** Xin Chen, Bin Hong, Zhangxi Lin, Jing Hou, ShunYa Lv, Zhendong Gao

## Abstract

The purpose of this study is to build a relatively complete human walking kinematic model. This model is combined with the rolling-foot model (lower limb) and multi-rod swinging model (upper limb) connected by COM. We calculated the velocity of COM and other critical joints of the upper limb by marker point capture experiment using the high-speed camera. This research shows that the hand joint velocity measured through the experiment can achieve high coincidence with that calculated by the theoretical model given specific inputs. Moreover, the common pattern of upper limb angles is also studied for an accurate description. The proposed kinematic model is expected to forecast desired motion intention for better compliance by the rehabilitation and assistive robots.

## 1. Introduction

The walking kinematic model is a profound simplification and generalization of the walking law of humans. The consummate walking kinematics model of human can provide theoretical guidance for gait planning of biped walking robot and lower limb dynamic exoskeleton, thus ensuring the robot obtain stable gait. The kinematic analysis of walking for the handicapped groups including hemiplegia and the design of the training track that conforms to their gait law is of great significance for the patients to recover their normal walking ability (Amatya, Sorkhabadi, & Wenlong, 2020; Liu, 2010; Wang, Yin, Yang, & Wang, 2018). Besides, the motion law originated from walking kinematic model is widely used in virtual human animation simulation (Cirio, Olivier, Marchal, & Pettre, 2013), human-machine interaction (Cifuentes, Bayon, Lerma, Frizera, & Rocon, 2016) and other fields. Nowadays, the difficulty of human walking kinematic model building is in that the human walking process includes instability and nonlinear characteristics. Therefore, it is not easy to describe the kinematic characteristics of human walking through a theoretical kinematic model.

The research focus of the human walking kinematic model primarily concentrates on the lower limb kinematic model. Kinematic models of lower limb walking mainly include first-order inverted pendulum model, second-order inverted pendulum model, spring inverted pendulum model, rolling foot model and so on. Tsai et al. (Kwon & Hodgins, 2010) created a dynamically balanced running character based on the inverted pendulum model. An et al. (Kang, Yingyuan, Yiran, Yunxia, & Chengju, 2018) incorporated joint torques and push-off impulse into the first-order inverted pendulum model to study the energy loss during walking. However, although this model can simplify the walking process to some extent, it is not natural and realistic for it assumes that the swing length of the inverted pendulum remains unchanged during the whole movement. Hu et al. (Jwu-Sheng, Kuan-Chun, & Chi-Yuan, 2013) compared the vertical fluctuation of COM under inverted pendulum and rolling foot model and then found that the vertical fluctuation of COM under inverted pendulum model was too large, revealing the shortcoming of inverted pendulum model. To describe the movement of lower limbs more accurately, the second-order inverted pendulum and the spring-damped inverted pendulum were proposed. Zhao et al. (Jianjun, Yi, Shihong, & Zhaoqi, 2014) synthesized human motion based on the second-order inverted pendulum to control the moment of hip joint and knee joint better. Compared with the inverted pendulum model, it shows more real lower limb motion from the geometric and physical point of view. Q.S. Yang (Yang, Qin, & Law, 2015) put forward the spring-damped inverted pendulum model and studied the influence of leg stiffness and angle of attack on the walking state. Actually, it can not describe the contact between the foot and the ground when walking.

Furthermore, a rocker-based inverted pendulum model, a rolling-foot model and a virtual-centre walking model were proposed to simulate the plantar curvature effectively during walking. Gard et al. (Gard & Childress, 2001) put forward the rocker inverted pendulum model and derived the walking parameters’ related equations. Xiang et al. (Qian, Hashimoto, Lei, Jianjun, & Shijie, 2019) improved this model by simplifying thighs and calves into rods and making ankle joints asymmetrically positioned on each foot, making the model more accurate. In addition, based on this model, they also derived a gait speed formula which is only determined by rolling factors. Hu et al. (Jwu-Sheng et al., 2013) estimated the COM velocity of human walking by using a wearable tri-axial accelerometer based on the kinematic model of the rolling foot.

Compared with the abundant kinematic models of lower limbs, scholars’ research on human upper limb kinematic models is relatively single, and the upper limb is usually simplified into several swinging rods for kinematics research. Hejrati et al. (Babak Hejrati, Chesebrough, Bo Foreman, Abbott, & Merryweather, 2016) modeled shoulder-angle and elbow-angle by Fourier series and then researched the effects of various conditions on arm-swing patterns during walking. Based on this model, Tanaka et al. (Yuki, Tomoya, & Yuji, 2018) proposed a dynamic model of arm-equipped rotational energy harvester (EH) during walking. Since the motion of arm swing is based on the movement of the lower limb, the combination of the upper and lower limb movements can effectively describe the movement of the arm. Further, the medium connecting the upper and lower body movement is precisely the COM of the body. As Hu et al. mentioned in their study, at present, most kinematic models focus on studying energy exchange mechanisms, force responses on the ground, metabolic rate, and so on. Nevertheless, few studies involved the use of kinematic models to estimate the velocity variation during walking of critical joints, including COM. Kinematic walking models, such as the connecting-rod model and rolling-foot model of the lower limb, can theoretically derive the displacement and velocity of critical joints including COM. The key aspect is to optimize the theoretical model with the experimental data.

In this paper, we propose an optimized kinematic model of human walking based on the kinematic model of the rolling-foot and multi-rod swinging model of the lower limb. The research point focuses on the corresponding relationship between the swing motion state of human upper limb joints (mainly hand joint) and lower limb gait characteristics (including step frequency and step length). The COM serves as the critical connecting point of the kinematic model of the upper and lower limb. We will estimate the COM velocity through the rolling-foot model, which is combined with data from the optical motion point capture experiment.

This paper is organized as follows. Section 2 presents the experimental layout, process and data processing of marker point coordinates captured by high-speed camera. Section 3 derives the velocity equation of joint points based on the rolling-foot model and multi-rod swinging model. Section 4 describes our proposed theoretical model of walking kinematics connected by COM. Section 4 evaluates the velocity of the hand joint measured in the experiment and that calculated by the theoretical model and shows the trajectory of critical joints. Section 5 discusses the paper with kinematical laws of critical angles and areas worthy of improvement in the future. Section 6 concludes the paper with the existing work and other ideas.

## 2. Methods

### 2.1. Participants

Six healthy subjects participated from a sample of individuals with healthy gait. Prior to participation, all participants provided written informed consent and the procedures of the study were approved by the university institutional review board. We used three male and three female subjects to account for the validity of the kinematic model for different gender. The age range of our subjects was 22-30 years (25.83±3.02 years) reported as (mean±standard deviation), and subjects’ body mass ranged from 48 to 81 kg (65.83±6.31 kg). The subjects’ height ranged from 1.58 to 1.8 m (1.70±0.05 m).

**Table 1.** Kinematic parameters of the upper and lower limb: Participant 1-6.

| Participant | Sex | Age(yrs) | Leg length(m) | Foot length(m) | Length from COM to<br>shoulder joint(m) | Length from shoulder<br>joint to elbow joint(m) | Length from elbow<br>joint to hand joint(m) |
| --- | --- | --- | --- | --- | --- | --- | --- |
| 1 | M | 23 | 0.94 | 0.28 | 0.44 | 0.28 | 0.28 |
| 2 | M | 27 | 0.98 | 0.29 | 0.48 | 0.295 | 0.3 |
| 3 | F | 29 | 1.05 | 0.285 | 0.47 | 0.32 | 0.32 |
| 4 | F | 22 | 0.98 | 0.245 | 0.43 | 0.31 | 0.3 |
| 5 | F | 24 | 0.97 | 0.24 | 0.45 | 0.34 | 0.28 |
| 6 | M | 30 | 1.05 | 0.29 | 0.45 | 0.33 | 0.32 |

### 2.2. Experimental set-up

As optical motion capture technology develops recently, the research and analysis of walking patterns have made significant progress. In optic-based motion capture systems, the human body is usually pasted with marker points (active type and passive type) at critical joints, then people walk within a limited range. Meanwhile, the equipped camera can carry out high-speed shooting and systematically track marker points in the walking process, to obtain the motion trajectory of walking, even other complex movements. Although such contactless measurement technology is relatively accurate, there exist some inherent problems: such systems are relatively expensive and difficult to operate, it requires relatively large fields and experimentation in specialized motion-capture laboratories. In this experiment, we adopted a method to capture human walking markers with a high-speed camera avoiding the partial problems. We used OpenCV to identify coordinate positions of critical markers in high-speed photos, then converted them into the actual distance for velocity calculation, so as to study the COM of human walking and the motion law of upper and lower limbs. The schematic diagram of the experimental layout is shown in Fig.1.

**Fig 1.**
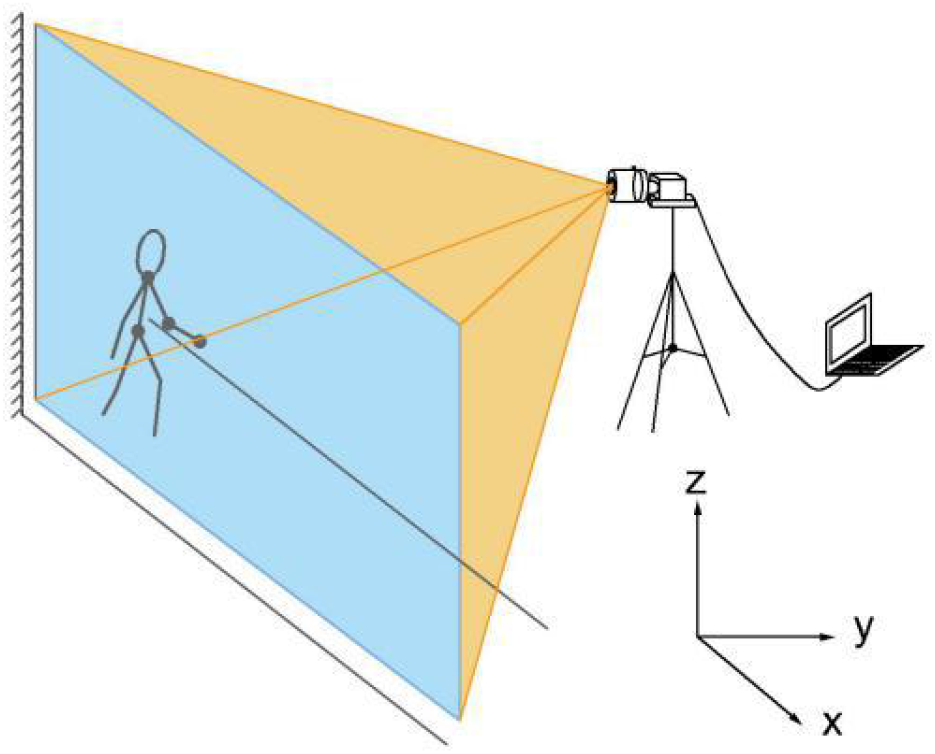
Experimental scene layout of the high-speed camera capturing marker points (Schematic diagram). The subject walked along the horizontal direction (x-axis), the z-axis represents the vertical direction and the y-axis represents the direction from the wall to the experimental equipment.

Firstly, the indoor experimental environment is arranged where the curtains are drawn and then the light brightness is adjusted to ensure that the room is in a relatively dim state, to facilitate the capture of luminous markers. Secondly, install the fixed-focus lens and tripod of the high-speed camera and keep the capture device level. The height of the tripod is adjusted to ensure that the subject’s entire walking process can be captured by the camera. In this experiment, the height of the tripod was set at about 90cm, the actual length of photos captured by the lens was about 5.1m, and the camera lens keeps perpendicular to the wall, shown in Fig.2. After the position of camera lens and tripod are fixed, the viewfinder frame will not be moved. The rate of photogeneration will have a particular influence on the trajectory capture and velocity calculation of luminous points. If the photogeneration speed is too slow, the accuracy of trajectory details will be affected, making the velocity calculation inaccurate. If the photogeneration speed is too fast, the small disturbance to the luminous markers will have a significant influence on the velocity calculation in a short time. It also puts forward higher requirements on computer performance. Due to these factors, the photo frame rate was set to 100fps on the MV Viewer, and a photo was saved every 1ms. Considering the limitation of reading and writing speed of computer hard disk, in this experiment, the interval time of generating two adjacent photos is 0.02s.

**Fig 2.**
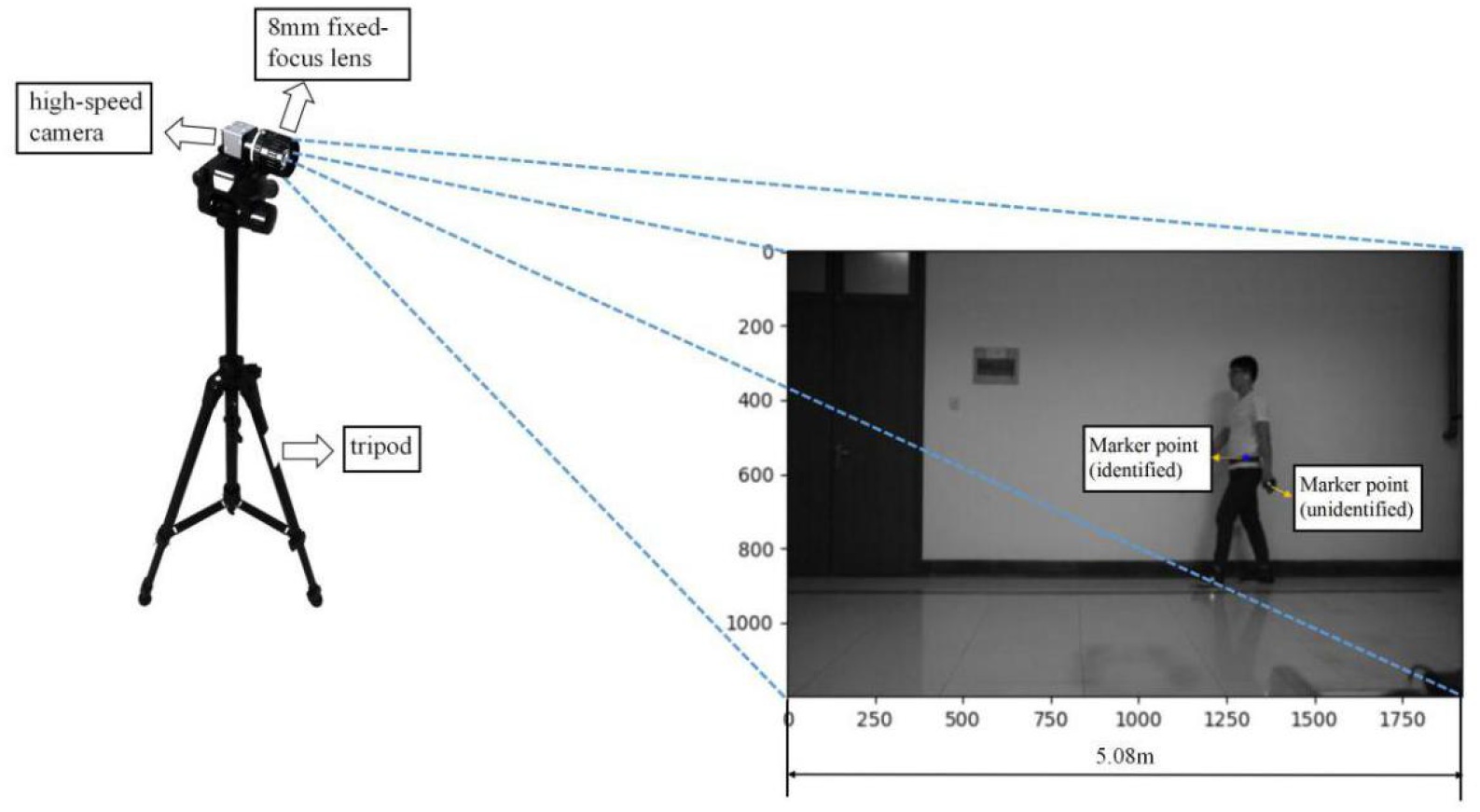
Experimental scene layout of the high-speed camera capturing Marker points (Experiment scene photo). Among them, experimental equipment includes the high-speed camera,8mm fixed-focus lens and tripod. The length of the photo frame is 5.08m, the unidentified marker point is white, while the identified marker point is blue.

### 2.3. Experimental procedures

As shown in Fig.2, a 94mm white light bulb was placed on the horizontal side of the subject’s critical joints, including the COM (approximately at the sacrum), shoulder, elbow, and hand joint. The white light bulb was fixed on the elastic belt with tape to avoid the error caused by a relative slide during walking. Meanwhile, according to the physical characteristics of each experimental object, the elastic belt was adjusted to a relatively tight state to prevent the elastic belt from moving up and down during walking. First, the experiment recorder was responsible for clicking the “continuous save” button of the upper computer software and starting the timing. Meanwhile, the experimental subject listened to the instruction of “start” and walked sideways on the wall at an average pace that was always parallel to the wall, keeping the whole walking process as natural and relaxed as possible. During the walk, the photos collected by the high-speed camera would be transmitted to the computer in real-time via the USB3.0 cable. When the hand marker was about to come out of the frame, the subjects stopped walking. Considering that the high-speed camera selected in the experiment had a delay time of about 1.2s after the end of shooting, the subject should stand in situ for about 1.2s after stopping walking, and the experiment recorder could press “stop saving”.

### 2.4 Data processing

After collecting the photos, it is essential to identify and locate the marker points. Here, we use the CVminmaxLoc() function in Opencv to capture the brightest marker point and Matplotlib to divide the coordinates of photos. Before the experiment, we converted the actual length of the reference object and the coordinate distance in the photo in equal proportion. Subsequently, we obtained that one coordinate unit in the photo is equal to the actual length of 0.002647m.

Inevitably, the marker point of the COM will be blocked by the swing of the arm, resulting in the loss of the coordinate information of the COM. The typical processing method is to complement the data loss by Lagrangian linear interpolation. In a complete walking cycle, the missing photo rate is about 5%-8%, this method is suitable for sample data supplement.

When calculating the average velocity of the marked points each time, it is often accompanied by a series of errors, including the inherent error of the instrument (the error of the photo interval), the random error, the error of the recognition time interval, and so on. In particular, due to the short interval between adjacent photos, the relative displacement error in the calculation caused by the coordinate error of the marker point will make the velocity calculation deviation larger. Here, we use the Kalman filter to eliminate the influence of data noises.

## 3. Theoretical Model of Walking Kinematics

### 3.1 Lower-limb kinematic model

To begin with, we aim to explore the relationship between COM velocity and step length, step frequency. From a macro point of view, the COM velocity of a pedestrian refers to the average velocity obtained by dividing the distance by the time.

However, the reality is that there exist slight differences in the frequency and length of each step, which results in the different velocity of COM in the walking process. Bao (Zhijun, Peisun, Jiangang, & Chunyu, 2000) and Zhang (Ye, Xiaojing, & Xiaotong, 2019) proved that the COM velocity during walking could be described by the curve of periodic fluctuation through the tracing point experiment and monocular video measurement experiment. However, their work only fitted the COM velocity curve with the sine function, yet the peak value of each wave was the same, which could not reflect the objective law of the fluctuation of the COM velocity when walking. We attend to use the rolling-foot model to describe COM’s motion: at each step of walking, the pressure applied to the ground moves forward from the heel to the toe. The integrated movements of ankles, feet and shoes during the support phase can be simplified into the rolling of the entire rotund rigid surface (Fig.3). Based on the rocker inverted pendulum model, related research had carried on the contact curve optimization between the supporting leg and the ground, then put forward the rolling-foot kinematic model of supporting leg movement. The earliest was presented by Gardetal (Gard & Childress, 2001). By means of the COM velocity formula of the rolling-foot model given by Hu, we calculated the COM velocity of walking. What’s different is that in our study, the step length and step frequency of each step obtained by the high-speed camera is substituted into the calculation, and the fluctuating COM velocity can be represented by step length and step frequency, so that more intuitive variables can be used to describe the state changes of people walking.

**Fig 3.**
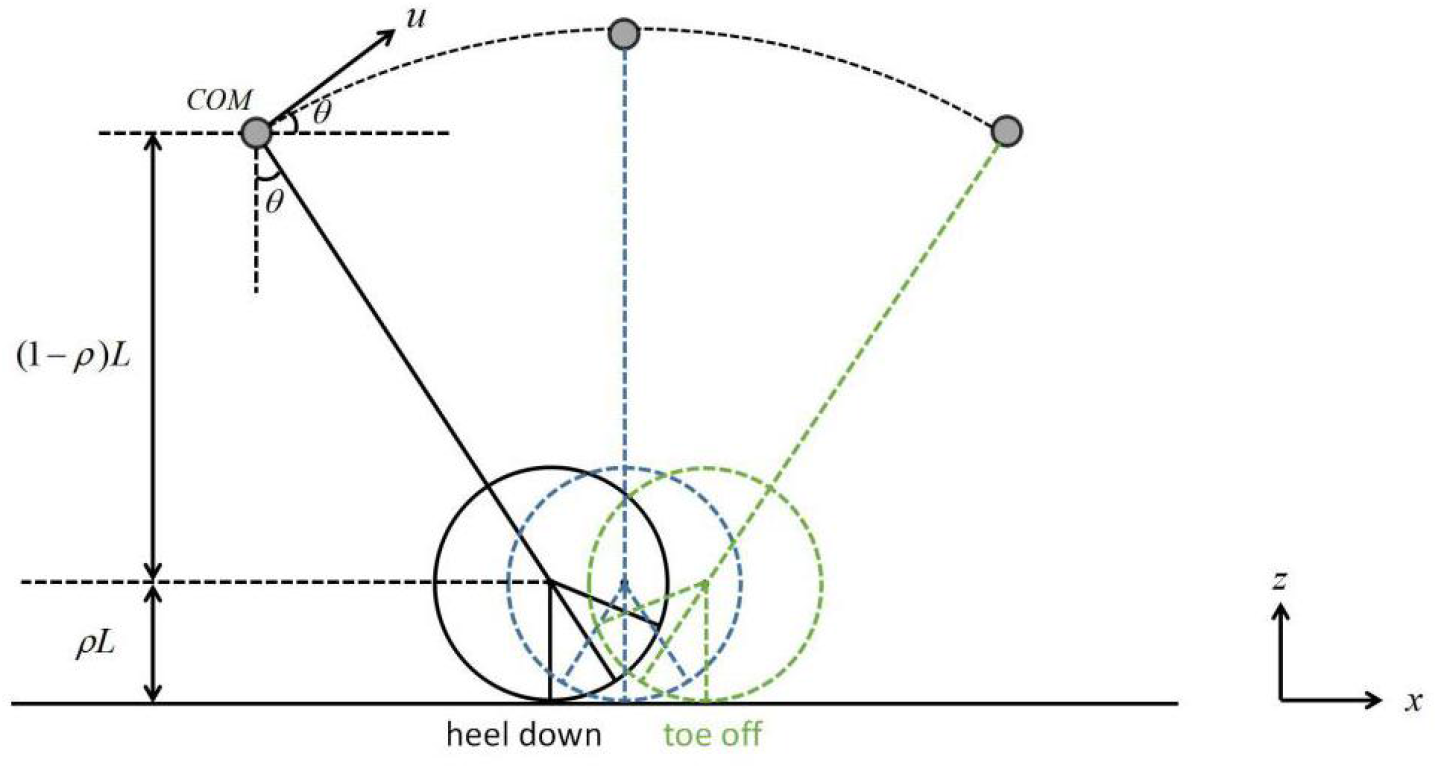
Rolling-foot walking model (Hu et al.,2013)

The centre of rotation is located above the ground contact point of a distance, which has a value of *ρL*, L is the leg length, and the rolling factor *ρ* varies from person to person (0 ≤ *ρ*≤1). The standing time T is defined as the duration between the point of contact (the heel-down) and the end of standing (the toe-off). Step frequency can be defined as the number of steps taken per time unit. The routine calculation is only to obtain the average cadence during a period of walking. However, as walking state changes, the step frequency also changes at any time. To accurately describe the variation of velocity in the walking process, our study took each step as a unit, measured the step frequency of each step, then established a relationship with the support period T in the rolling-foot model.

As depicted in Fig.4 and Fig.5, S1-S5 corresponds to 5 different states in the walking process, where the state marked the same number represents the same walking state. The walking state corresponding to the left leg served as a support leg during the standing period T is S1-S4, while S4-S5 is the transition process from support to swing of the left support leg, and the time experienced in this process is recorded as the swing time of the leg called t_swing_ (about 0.3-0.5s in normal walking). Meanwhile, during state S1-S5, the right leg also experiences a complete swing phase from the toe-off of state S2 to the heel-down of state S3. Thus, it experienced two steps (one on each leg) in state S1-S5. Set the time spent for one step t, length frequency *f* =1/ *t*, then the time containing two complete steps equals the support time t and the sum of the swing time t_swing_, namely:

**Fig 4.**
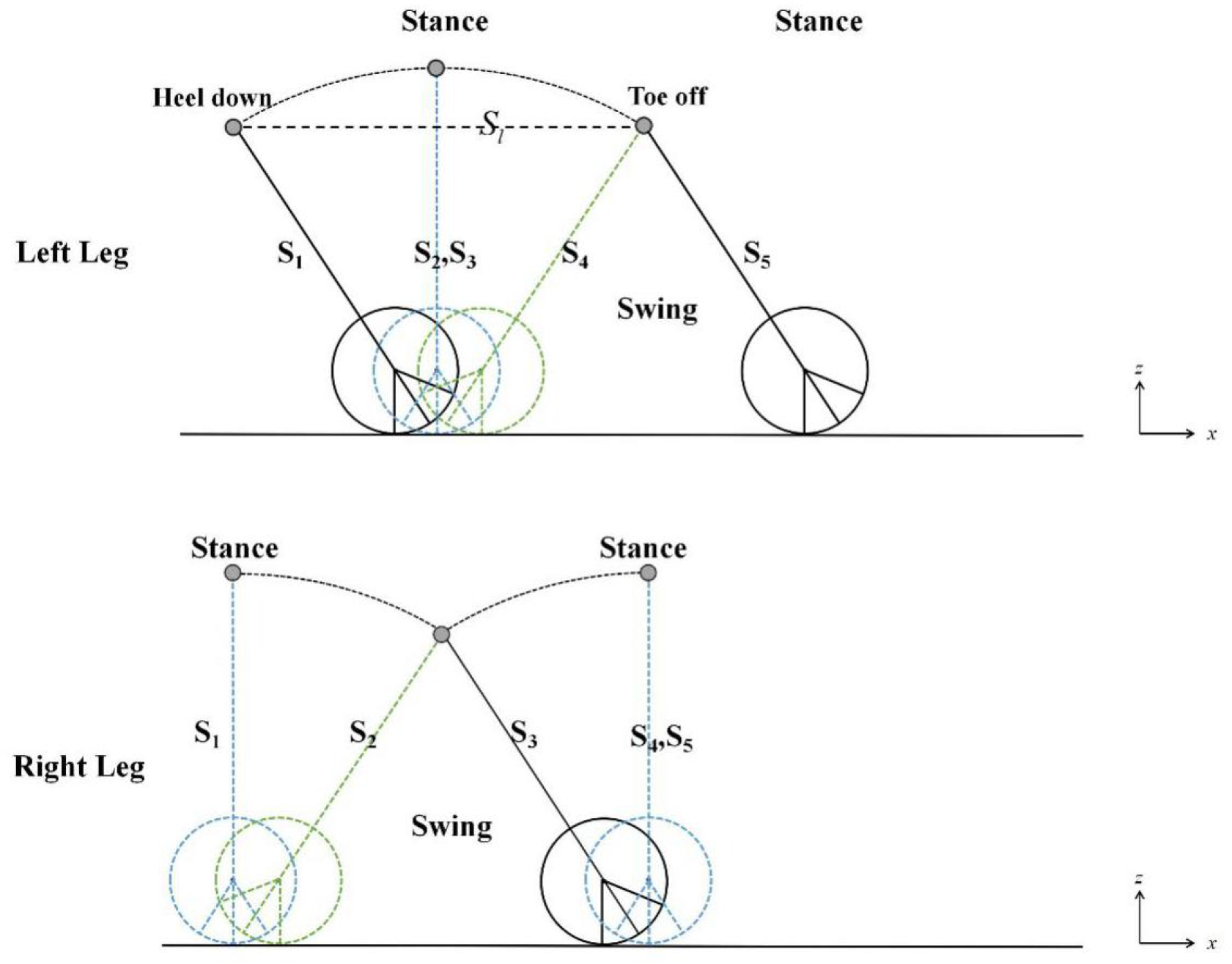
The alternating process of the support and swing of legs during a walking cycle (separate the left and right leg)

**Fig 5.**
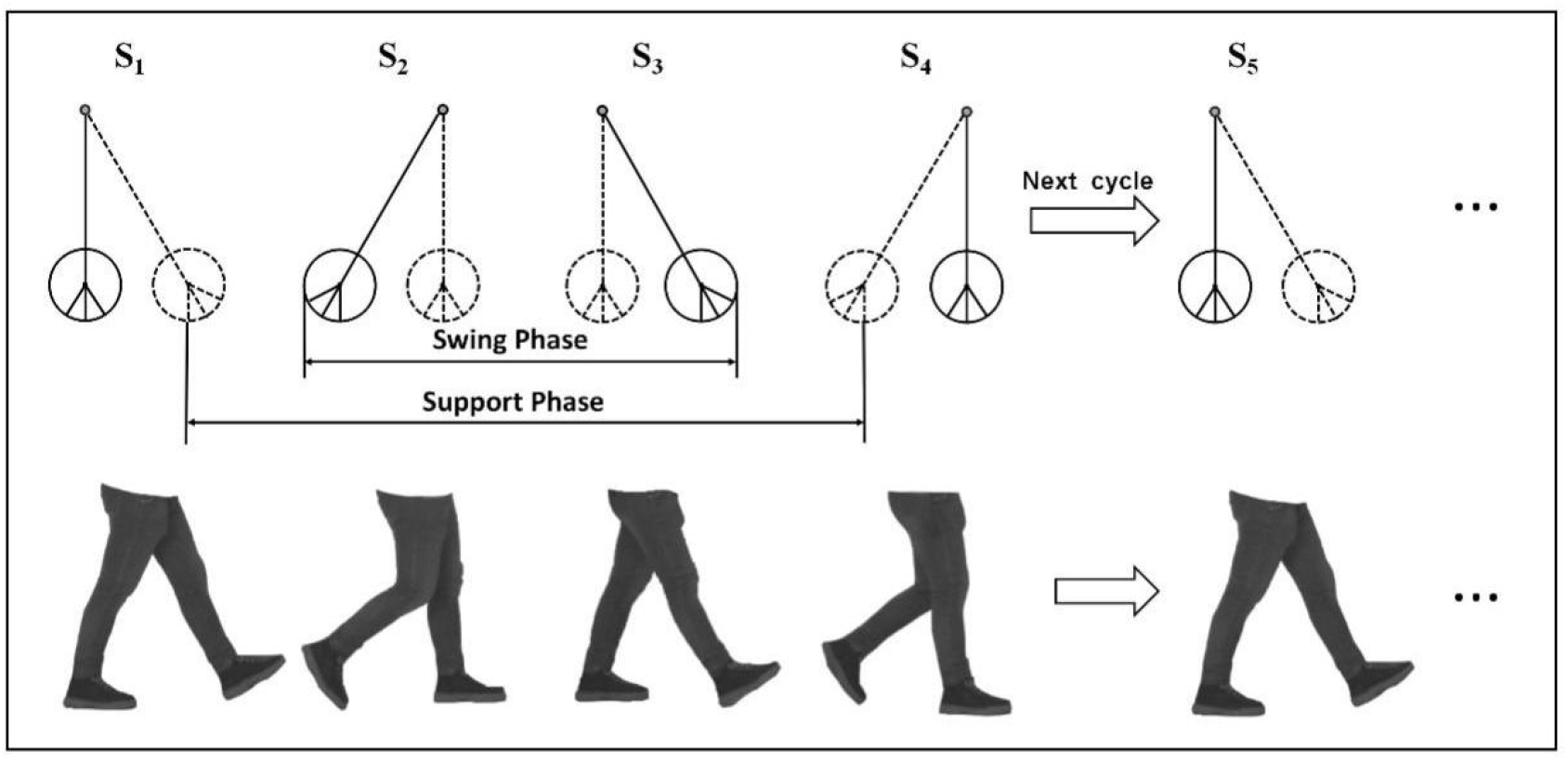
A walking cycle from heel-down of the left leg to next heel-down of the left leg. The figure below shows the actual walking condition of legs, which is consistent with the above figure.

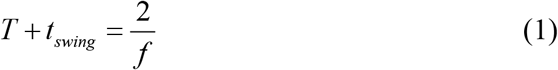

The step frequency can be expressed as:

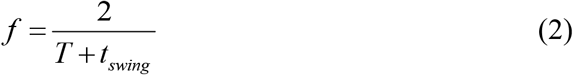

Angle α is defined as the maximum swing angle between the support leg and the vertical direction. It is assumed that angle α is the same at the starting point and endpoint of contact. During the swing of the inverted pendulum, the support leg remains rigid, the rolling-foot radius is *ρL*, and the length from COM to the centre of the rolling-foot is (1−*ρ*)*L*. Set the length of the foot to b, and then the following relationship can be established according to the geometric relationship in Fig.3:

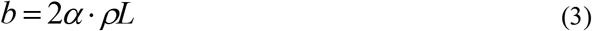

Since the angle α can hardly change in a stable walking state, the step length can be approximately simplified as the horizontal displacement from the initial point of the heel down to COM when the toe-off, that is to say, the horizontal displacement of COM is used to estimate the step length (marked *S*w in Fig.4). In practice, this estimation is based on the fact that the angle α during the adjacent walking period does not change much. If the adjacent period α varies greatly, this approximation can not be applied. Generally speaking, for the stable walking state, the step length can be approximately expressed as follows:

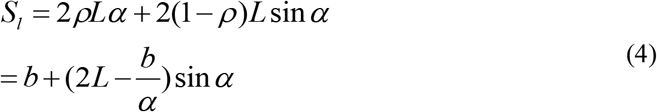

Assuming that the support motion of the support leg is a uniform process, the average tangential velocity of the COM under the rolling-foot model can be approximate as follows:

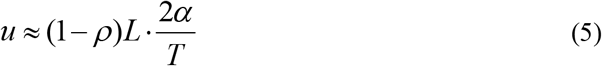

The length of the support leg is set to L, then the rolling foot radius is set to *ρL*.

The angle *θ*(*t*) is the swing angle between support leg and vertical direction at time t,*θ*∈[0,*α*]. The horizontal velocity of the COM can be regarded as the projection of the approximate tangential velocity of COM in the x-axis direction plus the horizontal velocity of the circle centre. We can get the velocity of COM as follows:

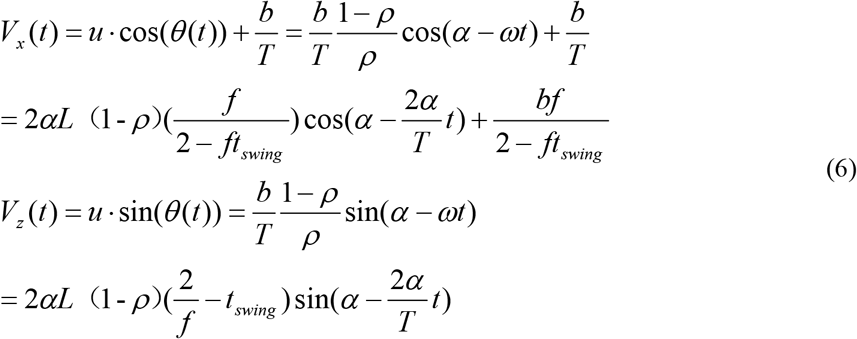

Among them, the maximum angle α between the support leg and the vertical direction has a corresponding function relationship with the step length *S*_l_ hence it is easy to get that the COM velocity can be described as a function related to the step length *s*_l_, step frequency f and time t. Both step length *s*_l_ and step frequency f show different values in each step, while t is a continuous variable.

### 3.2 Upper-limb kinematic model

Similar to the multi-rod swinging model of the lower limb, the upper limb model can also be simplified into a rigid multi-rod swing model (Fig.6). The angles between the upper body, upper arm, forearm and the vertical direction are respectively recorded as*θ*_1_, *θ*_2_, *θ*_3_. From the geometric relation, we can get that:

**Fig 6.**
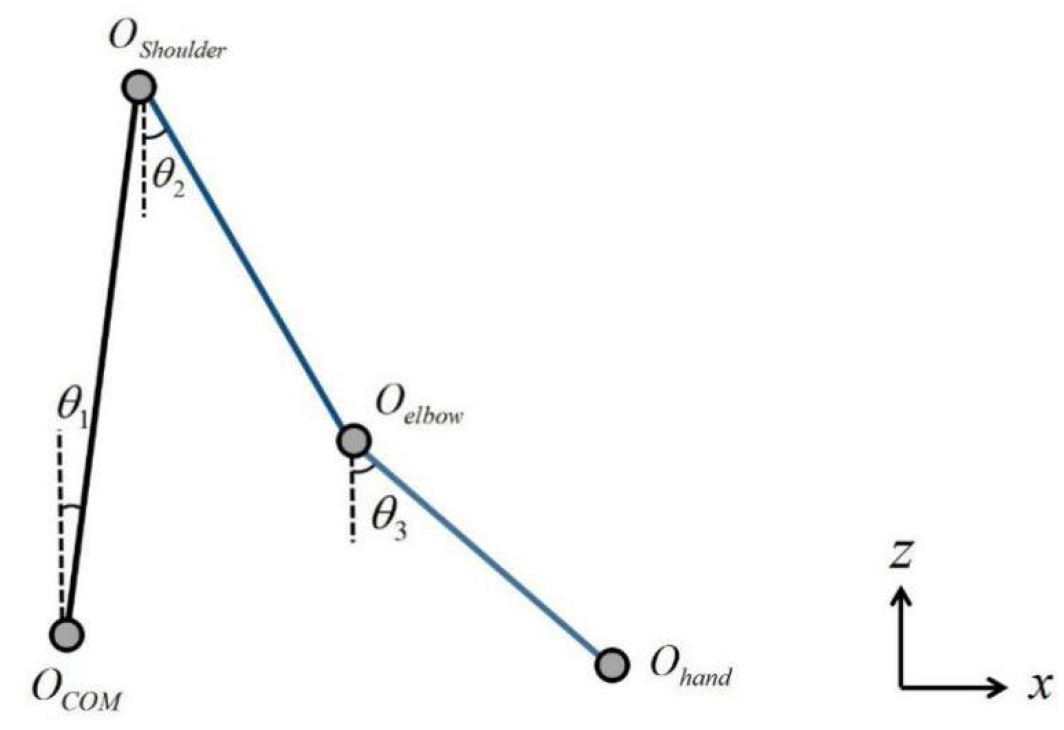
Multi-rod model of upper limb

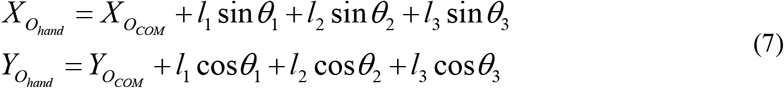

The derivation of both sides can calculate the swing velocity of the hand joint, it can be expressed as follows:

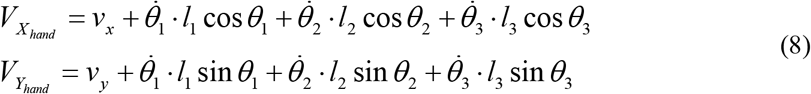

From the above kinematics expression of hand joint, it is easy to conclude that for a person with a certain height, the length of the upper limb, forearm and upper arm can be obtained from the empirical formula of human upper limb parameters. Hence the swing velocity of the human hand joint during walking is related to the COM velocity, add with three angles between each connecting rod and vertical direction. Since the COM velocity can be obtained by the rolling-foot model, we will continue to study the angle variation of several joints during walking.

### 3.3 Proposed kinematics model combined with upper and lower limb movements

Nowadays, most human kinematics models concentrate on the upper and lower limb separately but ignore the overall grasp of the human kinematic model. To fill this gap, we propose a human walking kinematic model with the combination of upper and lower limb movements, where the COM serves as the connection point of upper and lower limb motion transmission. The establishment of a complete kinematic model of human walking provides a sufficient theoretical basis for studying the kinematic law of the upper limb and exploring the motion relationship between the upper and lower limb.

As shown in Fig.7, the hand joint’s velocity is related to the step length and step frequency on the one hand, and the swing of the upper limb parts on the other. For a specific person in different walking states, the input of the upper and lower limb tends to be different, so the movement of the hand joint shows various characteristics. Through walking experiments, we have studied the changes in the input of the critical angles of the upper limb under normal walking conditions and modified the theoretical model based on the reliability of the experimental data.

**Fig 7.**
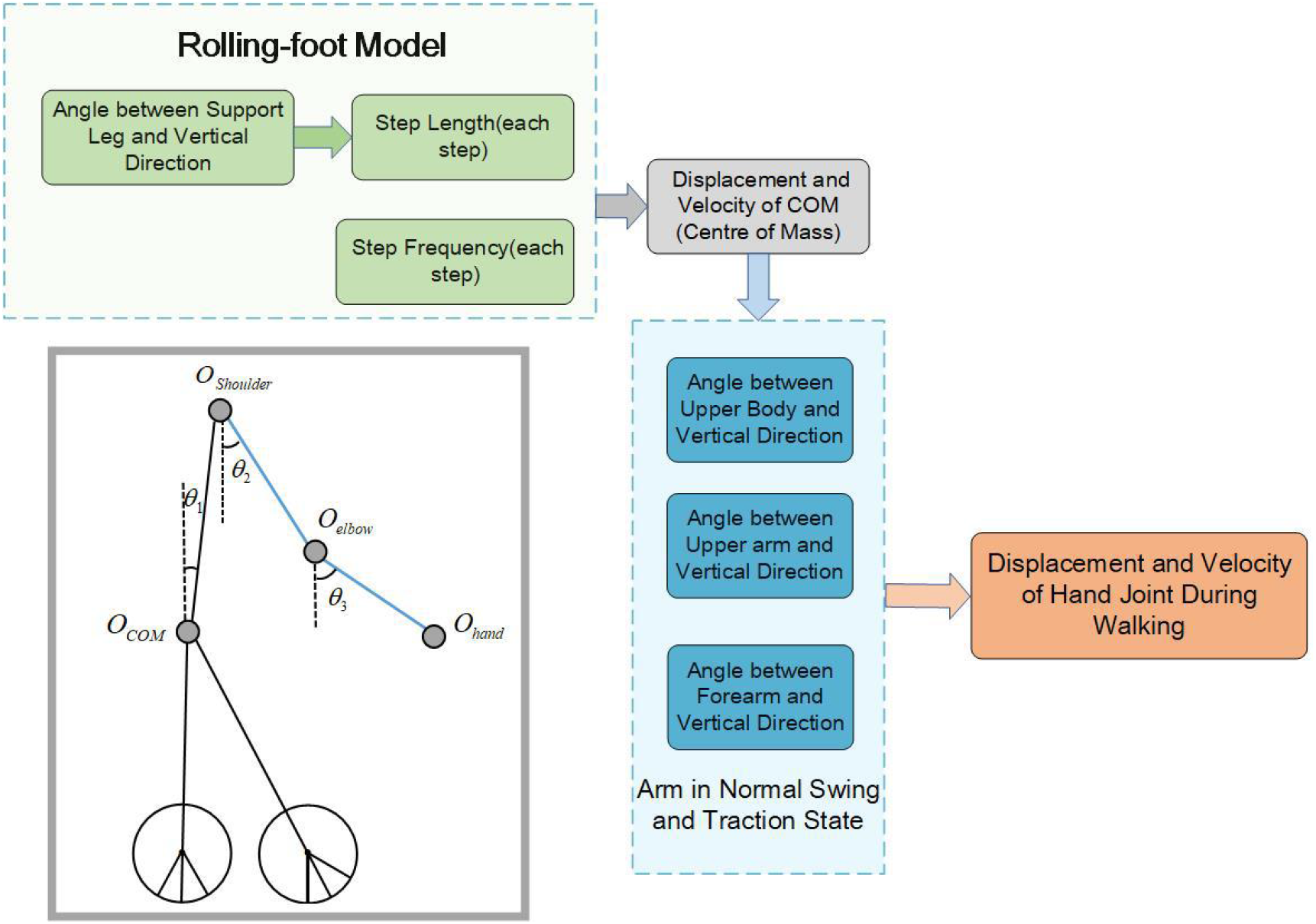
Human kinematic model combined with upper and lower limb movements

## 4. Results

### 4.1 Velocity of COM

To calculate the COM velocity based on the rolling-foot model, in addition to determining lower limb parameters (including leg length L and foot length b), input such as step length and support time should also be given. The step length and support time can be obtained from the photo data directly, while the angle α is obtained by the inverse solution of formula 4. We have tried to calculate angle α by connecting the COM with the mark points of the left and right foot joints, but the effect is not good. The angle α formed by left leg and the right leg differs by several degrees. The main reason is that the COM marker can’t be accurately pasted on the middle of the body.

As presented in Fig.8, in the general trend, the COM horizontal velocity calculated by the rolling-foot model is almost consistent with the COM horizontal velocity calculated by the marker point capture. Besides, it can also reflect the changing trend of the COM velocity showing an orderly fluctuation during the movement. We can find that in the velocity curve of COM, the peak height of each velocity wave is slightly different, which reflects the subtle change in velocity during walking. Since the experimental data are denoised by the Kalman filter, the actual experimental curve (red line) has a certain delay (about 0.5s) compared with the real situation. Take the velocity curve of subject 1 as an example, the experimental curve shows a total of six obvious waves, corresponding to six waves of the simulation curve. Besides, the relative height of the six peak waves of the experimental curve is almost consistent with that of the simulation curve. In fact, the peak value of the simulation curve wave is related to the angular velocity of the support leg, which can be expressed as *ωL*.

**Fig 8.**
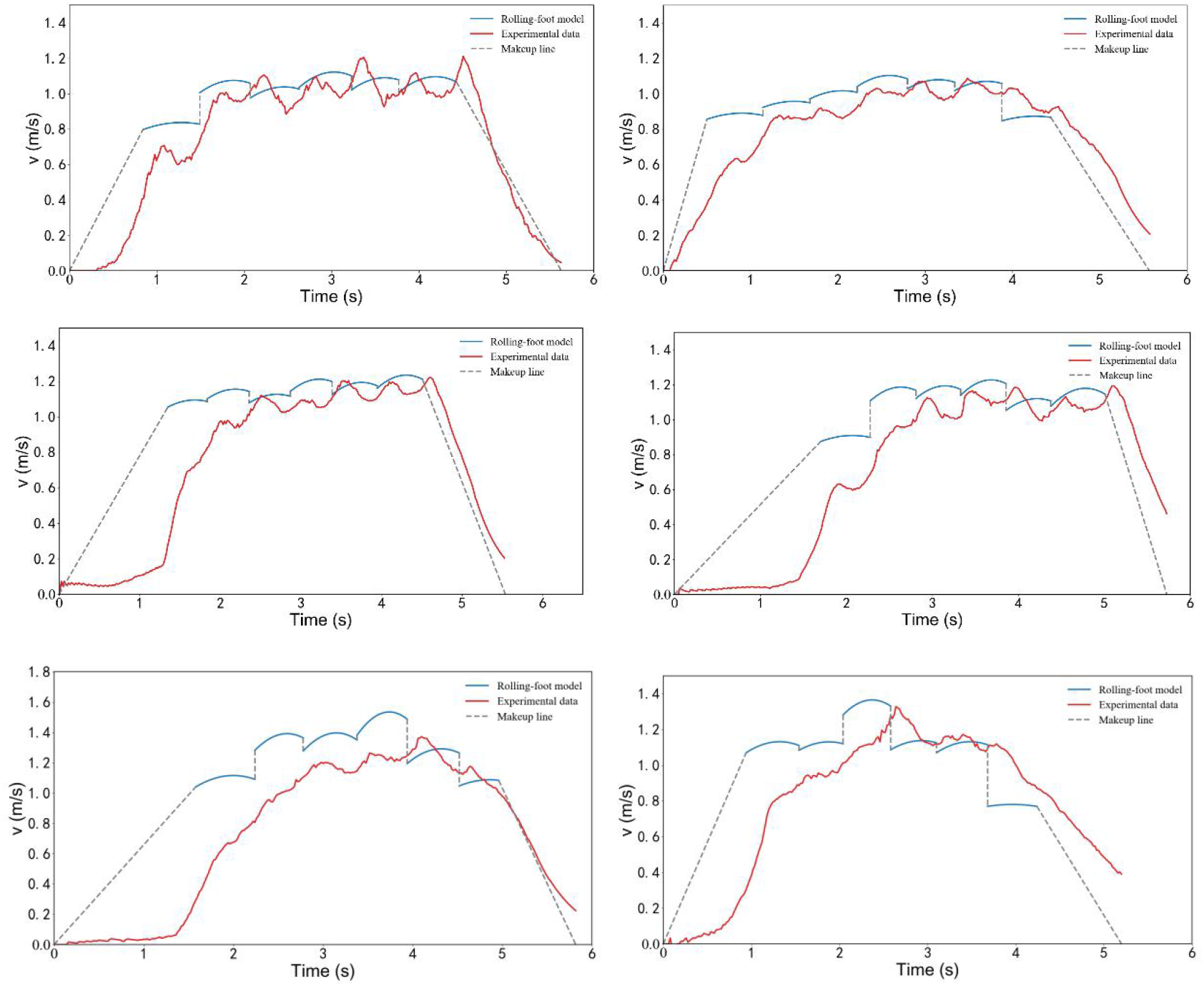
The velocity of COM in the horizontal direction of 6 subjects (blue curve: calculated by rolling foot model; red curve: obtained from experimental data (after filtering); gray line: as a supplement to the missing part of the curve, it has no data value)

The COM velocity calculated by the rolling-foot model is in steps, in other words, each step is packed with its corresponding velocity curve. In this way, since the number of steps is a discontinuous variable, the rolling-foot model can not solve the transition problem between adjacent steps, resulting in a trouble transition between adjacent steps. Besides, only when the walking state is relatively stable, the model can bring accurate velocity description. The model can not perform well when starting and stopping for the velocity expressed by the cosine curve is hard to change sharply in a short time. From Fig.8, it not hard to see that the first and the last wave curve can not perform well in describing velocity.

We find that the value of most velocity waves calculated by the model do not change significantly, and the difference between the maximum velocity and the minimum velocity in the same velocity wave is quite small, that is, the waveform is flat. The main reason is that the swing of the support leg is approximated to a uniform motion. Theoretically speaking, the swing of the support leg is a process of accelerating first and then decelerating. Therefore, the starting and ending velocities of the curve should be smaller than those of the existing model, while the peak velocity should be larger than those of the existing model. Here, we suggest that the angular velocity in the velocity equation be multiplied by a correction factor to better describe the change of angular velocity, so as to improve this situation.

Regretfully, due to the limitation of the laboratory space, the velocity curve can not be extended for better effect. From the existing curve, we can conclude that the calculated COM velocity in the stable walking state is relatively accurate.

### 4.2 Trajectory and velocity of upper limb critical joints

Six subjects were selected for the walking experiment and the whole walking process was kept natural and relaxed. The motion trajectory of the upper limb joint and the experimental photo was stitched together to make a moving map. Fig.9 shows the motion trajectory of each joint during the whole walking process (among which, the partially occluded COM coordinates were predicted), while Fig.10 depicts the walking images and joint trajectories of the last frame. It can be observed from the following figures that, first of all, the COM, shoulder, elbow and hand joint of the upper limb fluctuate regularly during stable walking. Among them, the motion trajectory of the hand joint is different from that of other joints: its motion trajectory presents a cyclic change trend of the slow rise and then rapid decline. The reason is that the swing velocity of the hand joint is mainly formed by the superposition of COM velocity and arm swing velocity. When the arm swings in the forward direction, the hand joint will be driven by the horizontal velocity of the COM and the trajectory will increase slowly with the arm swing up, and vice versa.

**Fig 9.**
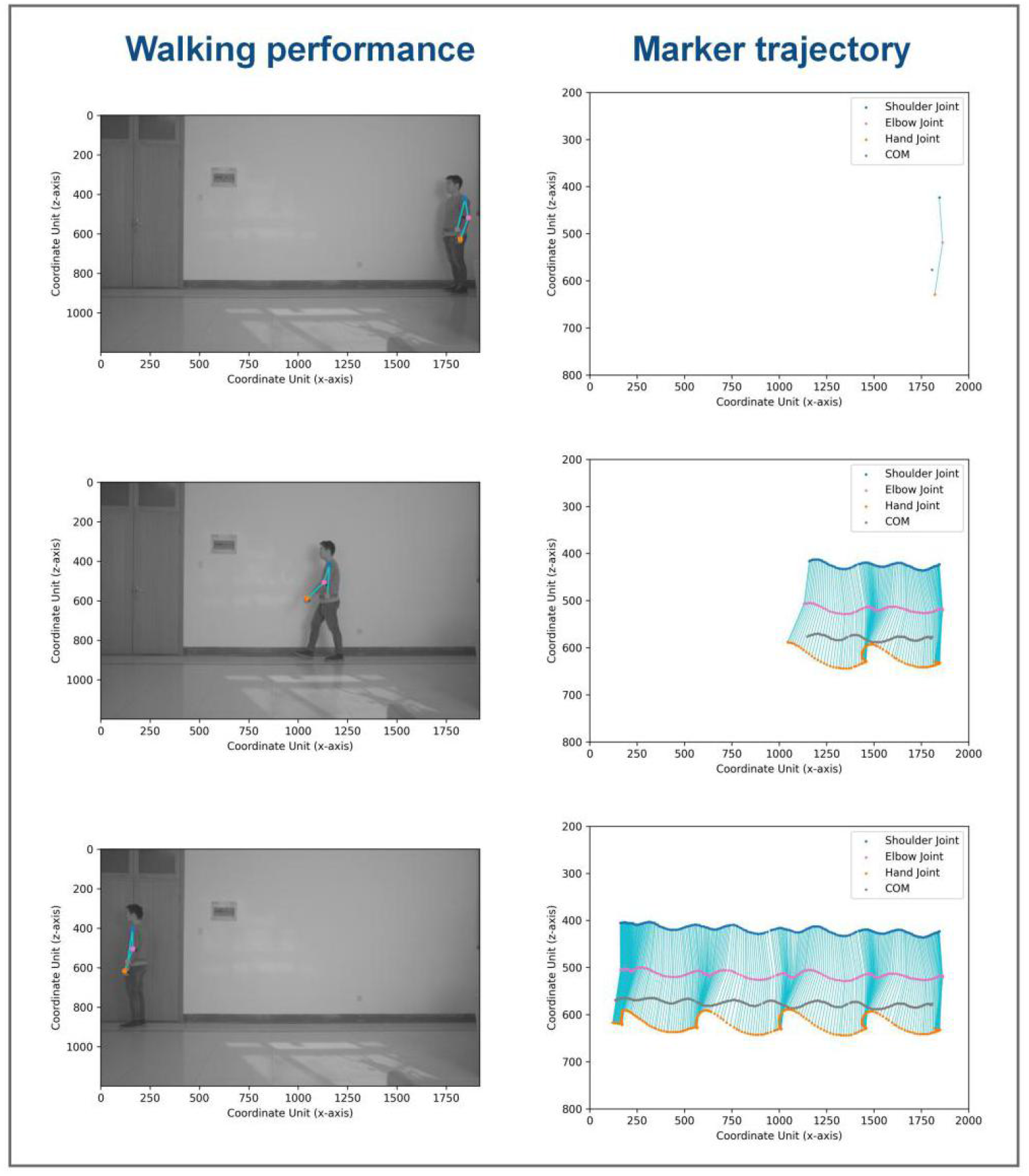
Mosaic of real-time walking images and joint trajectories (First frame, Intermediate frame and last frame) of Subject 1. From top to bottom, the trajectory of the shoulder joint, elbow joint and hand joint is represented by the blue line, pink line, gray line and orange line in turn. Walking direction: from right to left.

**Fig 10.**
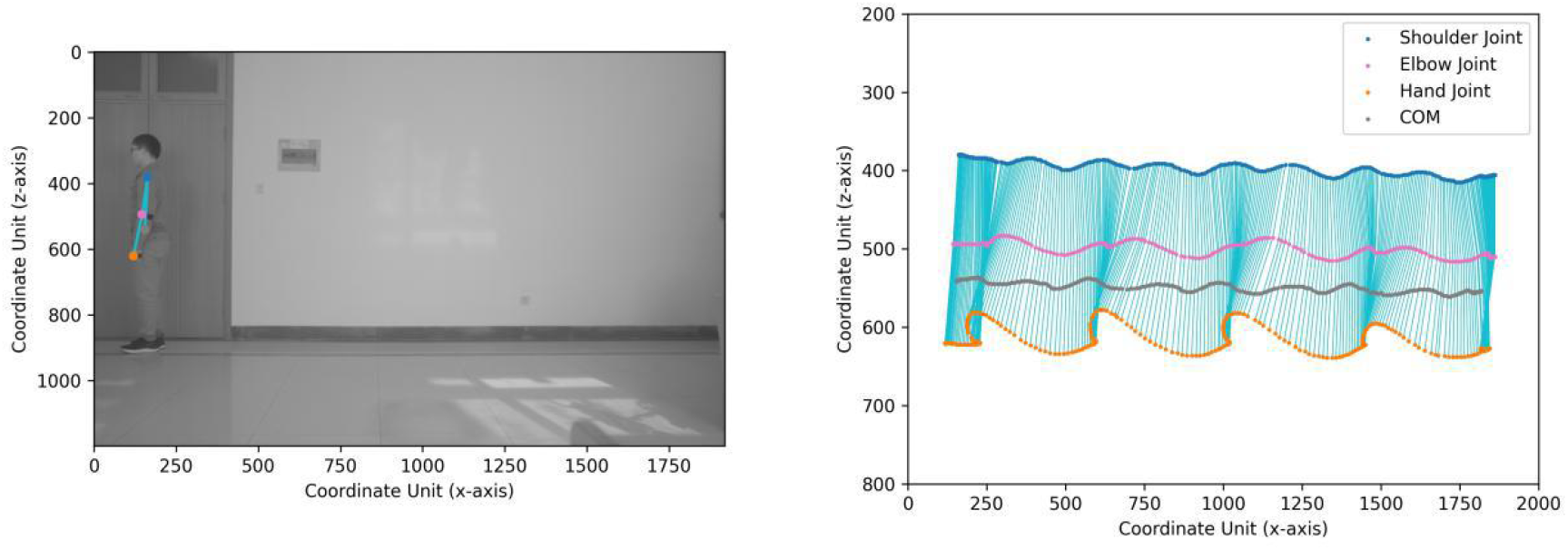
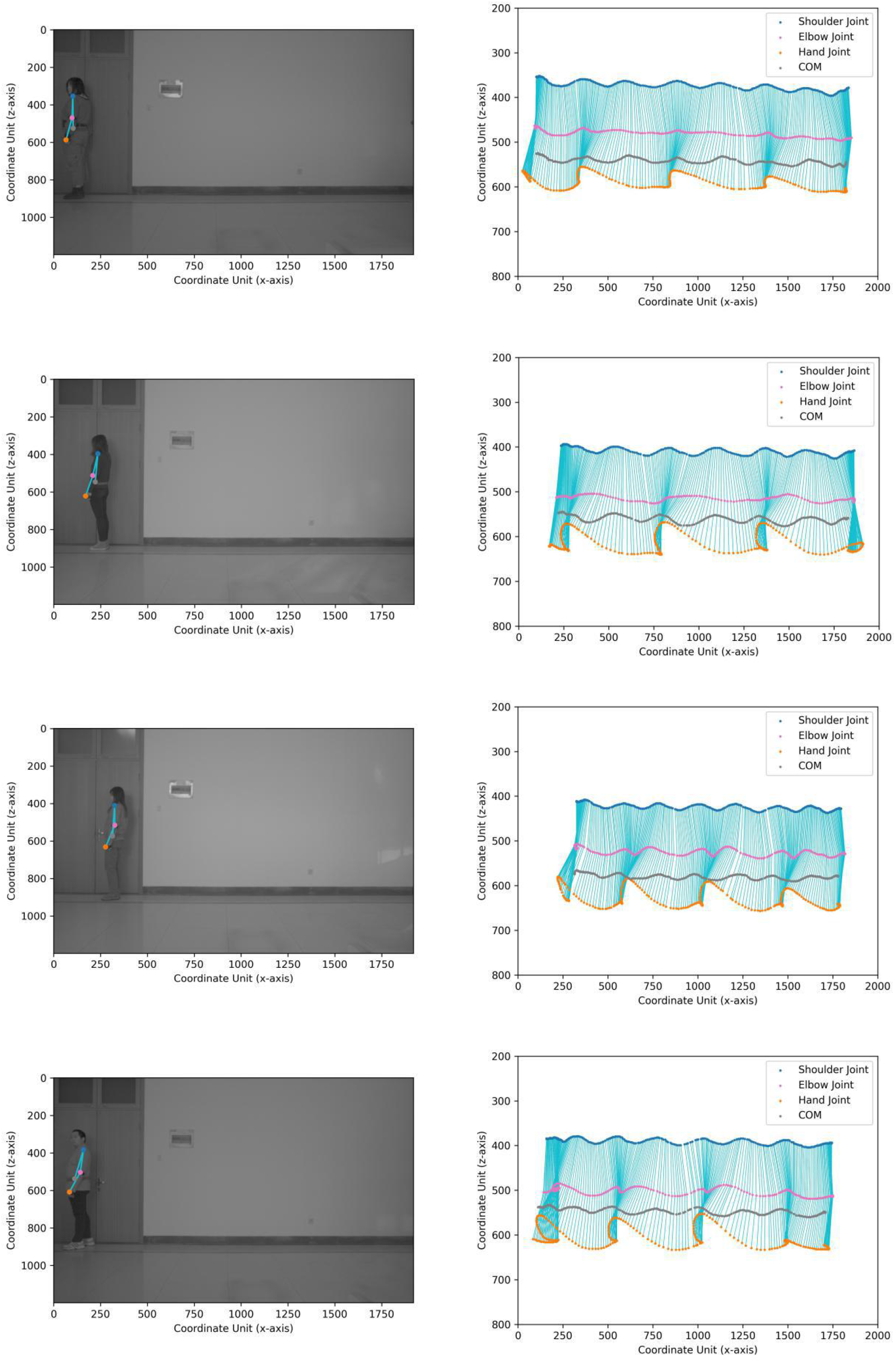
Mosaic of real-time walking images and joint trajectories (Last frame) of subject 2-6. The color representation of joint motion trajectory is the same as that in Fig.9.

Besides, it is not difficult to find that the blue line (connecting shoulder joint and elbow joint, connecting elbow joint and hand joint) is relatively dense when the hand joint reaches the highest point. The phenomenon reflects that the velocity of the hand joint is relatively slow when swinging to the highest point. The fluctuate range of the movement in the horizontal direction of the COM is small compared with the hand joint,and the horizontal movement of the hand joint can be decomposed into the arm swing (forward or backward) and the horizontal velocity of the COM. Therefore, in a certain frame of the whole process, the position of the hand joint is sometimes in front of the horizontal position of the COM, and sometimes after the horizontal position of the COM.

### 4.3 Validation of upper limb kinematic model

We have compared the hand joint velocity of the upper limb kinematic model under the input of known angle with that measured by the experiment. As shown in Fig. 11, the horizontal velocity of hand joint from the experiment and kinematic model can achieve high coincidence, while the vertical velocity shows subtle differences. From my view, the subtle differences of the vertical velocity can be owing to the effect of marker point capture. The vertical displacement of the COM and hand joint itself is quite small, a few coordinate units error will impose a certain influence on the velocity calculation. In general, it has been verified that the upper limb kinematic model can accurately describe the velocity of the hand joint when the input conditions such as model angle and angular velocity are given.

**Fig 11.**
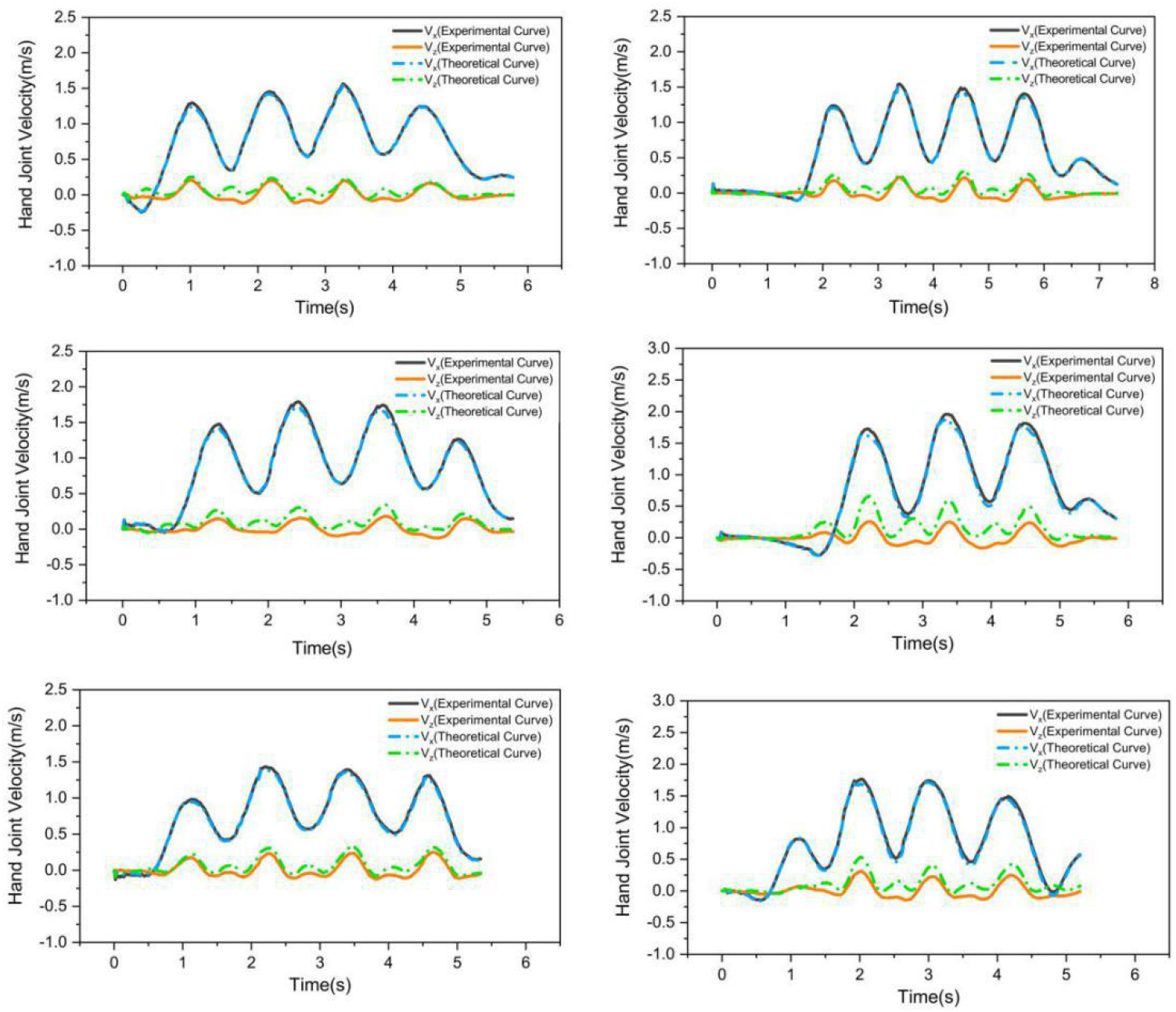
Horizontal and vertical velocity of hand joints from experimental measurement and theoretical model (Subject 1-6).

Meanwhile, we have noticed that the horizontal velocity of the hand joint is usually less than 0 at the beginning and end. When starting and stopping walking, the horizontal velocity forward of COM is relatively low, while that of the arm swing back is high in contrast, which results in the negative velocity of the hand joint in an absolute coordinate system. Some other laws of hand joint can also be obtained from Fig.11, for example, the horizontal and vertical velocity change synchronously more or less no matter increase or decrease, the velocity curve of hand joint can be fitted by a sine function, etc.

## 5. Discussion

In our research, the velocity of the hand joint is related to the swing of the upper body limb tightly. Therefore, studying the swing angle fluctuation of the upper body is of great significance for determining the movement of the hand joint. Different from most research that used IMU to measure angular velocities of human body rods (Babak Hejrati et al., 2016; B. Hejrati, Merryweather, & Abbott, 2018; Shangjie et al., 2018), we explored the movement of upper limb joint angle during normal walking based on the marker point capture experiment. Here, we set the joint coordinates of COM, shoulder, elbow and hand as (x_COM,_y_COM_), (x_shoulder,_y_shoulder_), (x_elbow,_y_elbow_) and (x_hand,_y_hand_), then the critical angle can be expressed as:

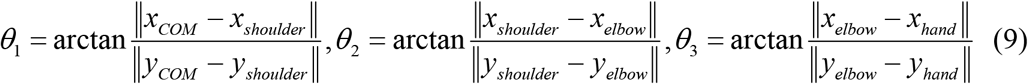

### 5.1 Joint angles of upper limb

In this model, we have defined *θ*_1_, *θ*_2_, *θ*_3_ as the input to describe the upper limb movement (Fig.6). Among them, *θ*_1_ refers to the angle between the upper body and vertical direction. Since each person’s body shape is different, it is difficult to determine the position of the COM at the same place, so the absolute value *θ*_1_ does not have much reference significance. However, the fluctuation of the *θ*_1_ curve shows that the upper body has a slight movement of forward and backward tilt in normal walking (Fig.12), and the difference between the upper and lower limits of *θ*_1_ can reflect the inclination degree of the upper limb during walking. From the experiment curve of six subjects, it reflects that *θ*_1_ shows a fluctuation range of about 10-20 degrees.

**Fig 12.**
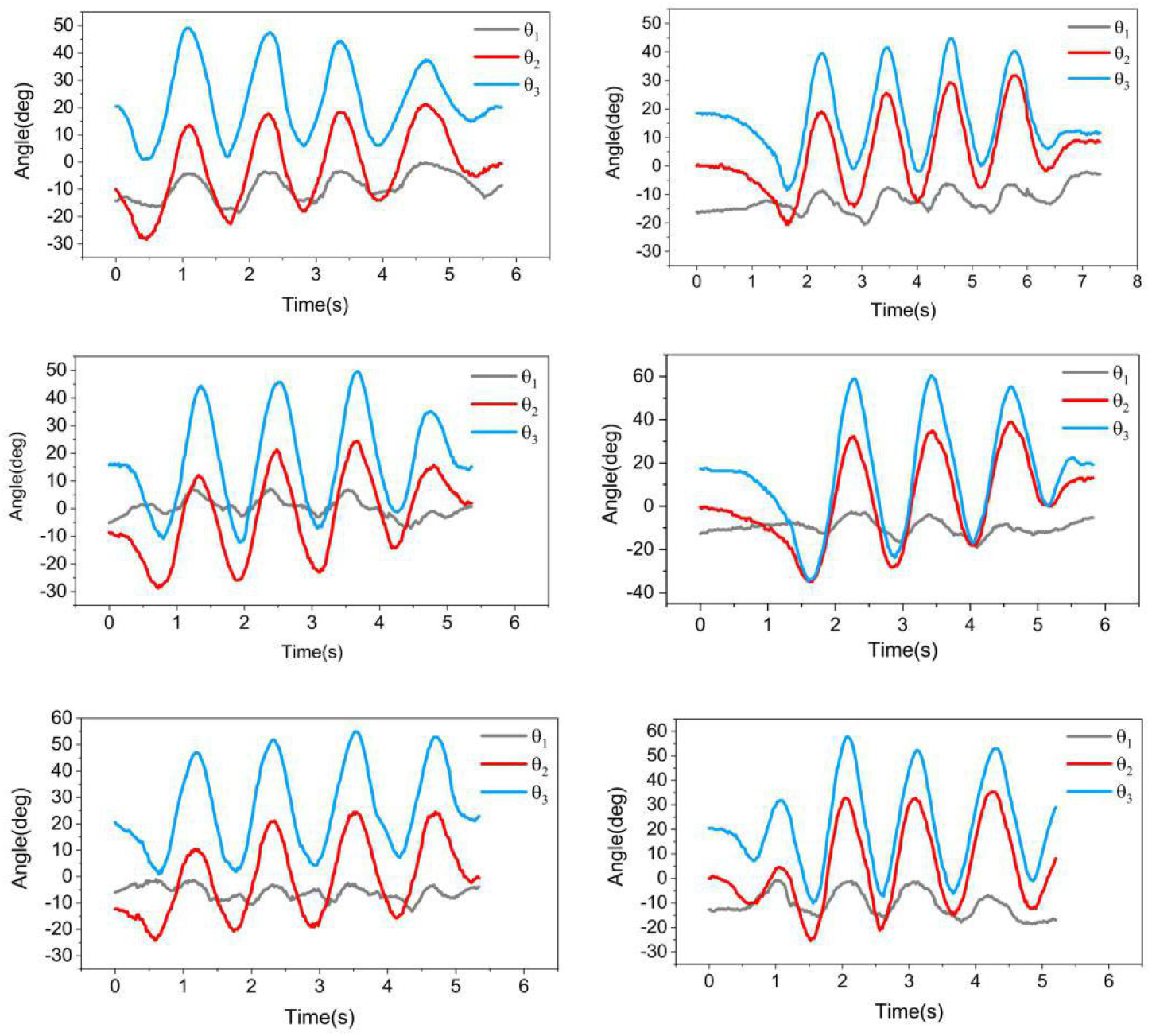
Joint angles of upper limb: *θ*_1,_ *θ*_2,_ *θ*_3,_ angle between the upper body and the vertical direction, angle between the upper arm and the vertical direction, angle between the forearm and the vertical direction)

It can also be found that the cycle of the three angles is roughly the same, in other words, the time coordinates corresponding to the peaks and troughs of the angles are quite similar. Tanaka et al.(Yuki, Tomoya, & Yuji, 2018) ever employed the equation for the elbow joint angle as the shoulder angle, it also verified the same cycle of shoulder joint and elbow joint. Hejrati et al. (B. Hejrati et al., 2018) generated an arm-swing trajectory while a subject increased and then decreased walking velocity, as the subject slowed, the peak of arm angle decreased. This law can be verified relatively obvious in Fig.12 of Subject 1,3 and 4 when slow to stop. For other subjects, the peak of the last angle before stopping doesn’t show an obvious drop mainly because the deceleration process is too short, which leads to the lack of deceleration transition process. In fact, it also exposed the shortcoming of this experiment, that is, limited imaging range leads to the lack of walking process continuity.

### 5.2 Joint angular velocity of upper limb

The angular velocity of each joint can be calculated by the differential of the measured joint angles. As shown in Fig. 13 the angular velocity of *θ*_2_, *θ*_3_is almost synchronous, and the absolute value of the upper and lower limits is almost slightly greater than that of the upper arm. The main reason is that the starting point of the swing arm is the same, when it reaches the highest position, the forearm will swing a little more than the upper arm due to the inertia. Besides, *θ*_1_ is also synchronized with *θ*_2_, *θ*_3_ in period, and the absolute value of angular velocity of *θ*_1_ has a certain reference value here, which reflects the inclination velocity of the upper body when walking. Similarly, its fluctuation period is approximately synchronized with *θ*_2_, *θ*_3_

**Fig 13.**
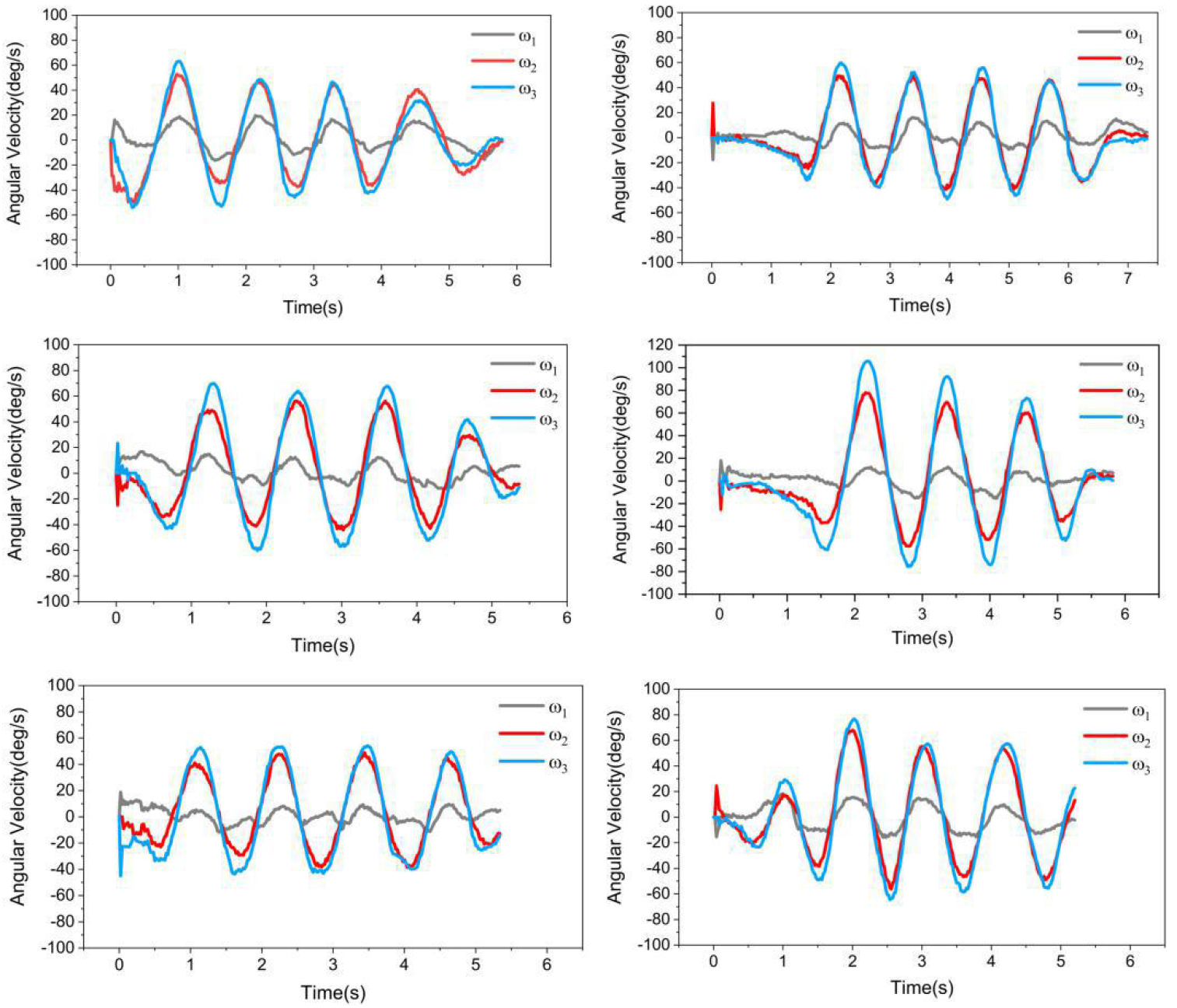
Joint angular velocity of upper limb: *ω*_1_, *ω*_3_, *ω*_3_

### 5.3 Reverse interpretation of the kinematic model

Now, the kinematic model of human walking has been established. Except for the kinematic parameters of the human body, the main inputs of the model are step length, step frequency and three critical angles of upper body, while the outputs are the horizontal and vertical velocity of hand joint. However, in practical application, it is rare to calculate the swing velocity of hand joint by inputting variables such as step length and step frequency. What we want to achieve is to judge the walking condition by the velocity of hand joint, which needs to swap the input and output of this model.

First, we discuss the inverse solution of the upper limb model. As shown in Fig.11, we can find that the horizontal velocity curve of hand joint is similar to the fluctuation of sine function, which is the result of the combination of COM motion and arm swing. Our goal is to theoretically infer the fluctuation of the COM and arm swing according to the velocity curve of the hand joint. In normal walking, the fluctuation of the horizontal velocity of the COM is smaller than that of the hand joint, which can be approximately uniform motion. When the COM velocity slows down, if the velocity of the arm swing remains unchanged, the velocity waveform of the hand joint will decrease as a whole, and the difference between the upper and lower vertices of the waveform is almost unchanged. But in most cases, after the COM velocity slows down, the arm swing velocity also slows down. The general velocity wave should show a trend of overall downward movement and smaller shape. This feature is more obvious in Subject 1, 2 and 4 in Fig.11 when the walking process stops. Similarly, as COM velocity increases, the arm swing velocity increases or decreases, there is a corresponding change law. Theoretically, it can preliminarily infer the changes of COM velocity and the velocity of the swinging arm.

When the joint angle model of the upper limb is given, the COM velocity can also be derived from equation (8). In the support phase of one step, the sum of the maximum and minimum velocity point is 2*b* / *T*, which the difference between the maximum and minimum velocity point is 2*b*(1− *ρ*) / *Tρ*. It means that the support time T and rolling factor *ρ* of this step can be calculated after the corresponding velocity waveform of one step is determined. According to equation (3) and (4), the step length corresponding to this step can also be obtained. Besides, there is a certain relationship between step frequency and support time T. Generally speaking, the processing and calculation of the COM horizontal velocity waveform can theoretically infer the state of people in the walking process. It is of great significance for judging the walking state from the perspective of intuitive parameters.

### 5.4 Additional opportunities and limitations

In this study, the walking kinematics model in theory mainly refers to the general model in the walking process, focusing on the corresponding relationship between the hand joint velocity and the state of the upper and lower limb. In fact, in the field of human upper limb kinematics, most researches are to provide theoretical guidance for the upper limb exoskeleton and other wearable robots, so as to carry out more suitable rehabilitation treatment for patients. Rosen et al. (Rosen, Perry, Manning, Burns, & Hannaford, 2005) collected the kinematic data of the human arm during daily activities and found that compared to a healthy operator, the exoskeleton might be utilized differently by a disabled person. Tang et al. (Shangjie et al., 2018) studied patiotemporal kinematics synergies on upper limb movements, which showed the differences existing in upper arm. In addition, the critical angular velocity of the upper limb can also be used for the desired motion estimation in assistive robots (Khan, Khan, & Han, 2016). Therefore, it is feasible to use the kinematic model to identity the human walking state and realize human-machine interaction.

In our research, our ultimate goal is to use the flexible cable to realize the interaction between visually impaired people and guide dog robot. Therefore, the key to the kinematics model of the people is the motion state of the hand, which will have a great impact on the configuration and force of the flexible cable. The proposed model of this study might be considered as the starting point for coupling people, flexible cable and guide dog robot dynamics model. Future works will extend the research on the upper limb kinematics when grasping a flexible cable.

Of course, this study inevitably ran into some limitations. First of all, in the experimental means, some limitations exist in the experimental field of motion capture. The limited walking space makes it difficult to observe the whole process of acceleration and deceleration. Our solution is to use six-axis IMU, for it can integrate to get COM velocity and solve the problem that the marked points are blocked by the arm. Meanwhile, the relevant angle can also be measured. In the description of the model, due to the high randomness and uncertainty of the human walking state, the description of the model in this study is not detailed enough. It is expected that in the future, the random and diverse characteristics of different people walking, especially disabled people whose gait is usually characterized by instability and heaviness, will be added to our proposed model for better performance of assistive robots.

## 6. Conclusion

Overall, the rolling-foot model can describe COM velocity in a steady walking state accurately, and the corresponding COM horizontal velocity of each step can be regarded as a function of step length, step frequency and time. Meanwhile, given the critical angle of the upper limb, it is feasible to simplify the upper limb into a multi-rod swinging model to calculate the velocity of the hand joint, further building the relationship between hand joint and lower limb movement situation. Based on these results, we combine the kinematic models of upper and lower limb and explore the input rules of some key angles. Of course, these angle inputs reflect the state of human walking to a certain extent, which shows various trends for people in different states (negative or positive mood and so on), is worthy of further study.

## Acknowledgments

The authors thank Xin Chen, Jing Hou, Zhangxi Lin, Ya Ji, Jiayue Qing, Hao Zhang, Chengwei Yang and Bin Zhao for their assistance in data collection.

